# Unique roles of ATAC and SAGA - KAT2A complexes in normal and malignant hematopoiesis

**DOI:** 10.1101/2020.05.14.096057

**Authors:** Liliana Arede, Elena Foerner, Selinde Wind, Rashmi Kulkarni, Ana Filipa Domingues, Svenja Kleinwaechter, Shikha Gupta, Elisabeth Scheer, Laszlo Tora, Cristina Pina

**Affiliations:** Department of Haematology, University of Cambridge, Cambridge, United Kingdom; Department of Genetics, University of Cambridge, Cambridge, United Kingdom; Institut de Génétique et de Biologie Moléculaire et Cellulaire, 67404 Illkirch, France; Centre National de la Recherche Scientifique (CNRS), UMR7104, 67404 Illkirch, France; Institut National de la Santé et de la Recherche Médicale (INSERM), U1258, 67404 Illkirch, France; College of Health and Life Sciences, Division of Biosciences, Brunel University London, Uxbridge UB8 3PH; Wellcome Trust Cambridge Stem Cell Institute and Department of Haematology, University of Cambridge, Cambridge, United Kingdom

**Author notes:** Contributed equally (Elena Foerner, Selinde Wind).

**Keywords:** ATAC, SAGA, Self-renewal, Differentiation, Leukemia, Normal hematopoiesis

## Abstract

Epigenetic histone modifiers are key players in cell fate decisions. Significant research has focused on their enzymatic activity, but less is known about the contextual role of the complexes they integrate. We focus on KAT2A, a histone acetyltransferase we recently associated with leukemia stem cell maintenance, and which participates in ATAC and SAGA complexes. We show that ATAC is uniquely required for maintenance of normal and leukemia stem and progenitor cells, while SAGA more specifically contributes to cell identity. This dichotomy sets a paradigm for investigating epigenetic activities in their macromolecular context and informs epigenetic regulator targeting for translational purposes.

## INTRODUCTION

Histone acetyltransferase (HAT) coactivator complexes are key determinants of cell fate decisions, facilitating gene expression through enzymatic and nonenzymatic activities (Sun et al. 2015). Extensive work has focused on the enzymatic effectors of HAT complexes, but the contributions of the macromolecular context in which they operate is less well characterized. ATAC (Ada-Two-A-Containing) and SAGA (Spt-Ada-Gcn5-Acetyltransferase) are two metazoan multisubunit complexes (Fig. 1A) that share the same HAT: KAT2A or KAT2B (Spedale et al. 2012). Integration of KAT2A in either complex is a requirement for its full acetyl-transferase activity (Riss et al. 2015). Moreover, the two complexes exhibit different chromatin specificities and regulate distinct sets of genes (Krebs et al. 2011). This highlights the role of complex composition and the interaction of the different subunits with KAT2A in directing its activity. KAT2A is the human orthologue of yeast Gcn5 and is required for H3K9 acetylation (H3K9ac) (Brownell et al. 1996). Moreover, KAT2A was recently shown to catalyse succinylation, another acyl-modification which, like H3K9ac, is associated with transcriptional activation (Wang et al. 2017a). Kat2a HAT activity is required for appropriate mouse mesodermal specification (Xu et al. 2000) and *Drosophila* oogenesis and metamorphosis (Carre et al. 2005). Additionally, Kat2a plays a role in maintaining stemness (Moris et al. 2018) and is essential for survival of neural stem and progenitor cells (Martinez-Cerdeno et al. 2012). Interestingly, Kat2a has not been implicated in stem cell maintenance in the hematopoietic system (Domingues et al. 2020), the most extensively characterized model of stem cell fate decisions (Jacobsen and Nerlov 2019). Hematopoietic stem cells can give rise to up to 14 different cell types by probabilistic restriction of lineage choices (Notta et al. 2016) with early separation of megakaryocytic (Carrelha et al. 2018) and erythroid fates (Belluschi et al. 2018), which precede resolution of myeloid and lymphoid lineages (Belluschi et al. 2018). Although not required in stem cells, Kat2a participates in myeloid and lymphoid cell differentiation, acting through histone (Kikuchi et al. 2014; Gao et al. 2017) and non-histone (Bararia et al. 2016; Wang et al. 2017b) protein acetylation control of gene transcription.

**Figure 1.**
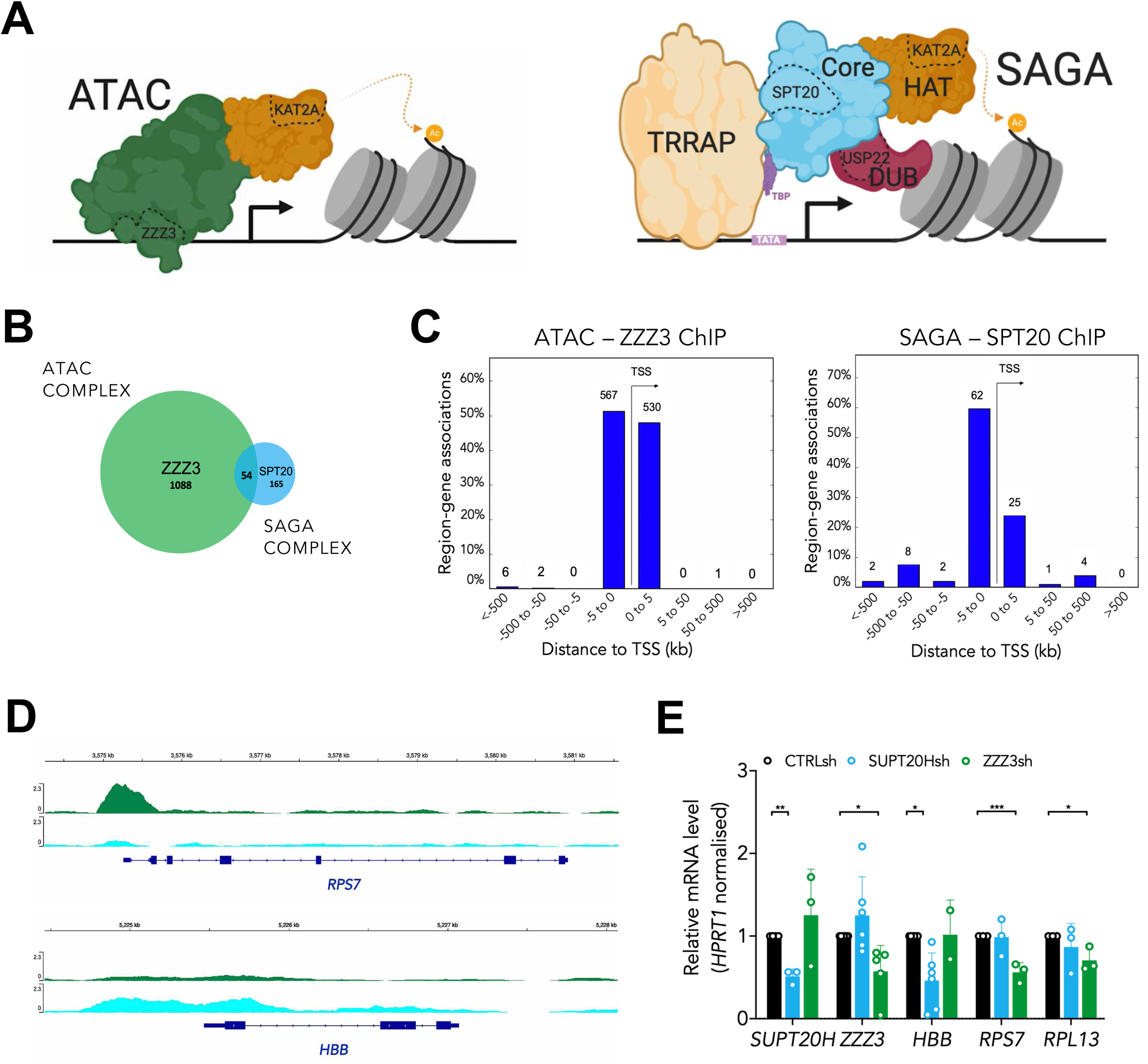
KAT2A-containing ATAC and SAGA complexes have unique targets in haematopoietic cells. **(A)** Schematic representation of the human ATAC (left) and SAGA (right) multiprotein complexes (subunits tested in this study are highlighted). **(B)** Venn-diagram of consensus ZZZ3 and SPT20 ChIP-seq binding from 2 independent experiments. **(C)** Genomic location of ZZZ3 (left) and SPT20 (right) ChIP-seq binding in K562 cells. Summary of consensus peaks from 2 independent ChIP-seq experiments is shown. **(D)** Representative ChIP-seq peak for ZZZ3 target in K562 cells (*RPS7*) and representative ChIP-seq peak for SPT20 target in K562 cells (*HBB*). **(E)** Quantitative RT-PCR analysis of expression of ATAC and SAGA complex targets in K562 cells. N≥3 independent experiments, mean ± SEM of gene expression relative to *CTRLsh*, normalised to *HPRT1* housekeeping gene. Two-tailed t-test for significance *p<0.05, **p<0.01, ***p<0.001.

The yeast KAT2A-containing SAGA was the first multimodular HAT complex to be isolated (Grant et al. 1997). Since then, several studies unveiled its molecular architecture: SAGA comprises 19 subunits organised in 4 functionally distinct modules (Liu et al. 2019), a structure highly conserved from yeast to man (Spedale et al. 2012). In addition to the KAT2A-participating HAT module, which also includes TADA2B, TADA3 and CCDC101 (SGF29), SAGA contains: (1) a H2B de-ubiquitination (DUB) module centered on USP22 enzymatic activity; (2) a Core module comprising SPT20, which is specific to SAGA, and 5 TATA-binding protein associated factors (TAFs), 3 of which are shared with TFIID; and (3) the transcription factor interaction module, TRRAP. Two recent reports on the Cryo-EM structure of the yeast SAGA complex described the interactions between the different modules with key implications for gene activation (Papai et al. 2020; Wang et al. 2020). The SAGA Core module contains an octamer-like fold that facilitates TBP loading onto TATA promoters (Papai et al. 2020); the two enzymatic HAT and DUB modules connect flexibly to the Core (Papai et al. 2020; Wang et al. 2020), suggesting functional independence between them. The ATAC complex, in turn, is exclusive to multicellular eukaryotes (Spedale et al. 2012). It was first linked to chromatin remodeling functions in *Drosophila* (Carré et al. 2008), where it preferentially targets histone H4 (Suganuma et al. 2008). ATAC histone substrates in mammalian cells are less clear, although human ATAC preferentially modifies H3 (Helmlinger and Tora 2017). ATAC specific elements include DNA-binding subunit ZZZ3, and YEATS2, which is required for assembly of the ATAC complex and has acetyl-reading activity (Mi et al. 2017). ATAC comprises an additional HAT activity by KAT14, which is essential for ATAC assembly and required in embryonic development (Guelman et al. 2009).

In addition to functions in normal development, both KAT2A-containing complexes have been implicated in malignant transformation (Koutelou et al. 2010). ATAC-*YEATS2* was shown to be highly amplified in non-small cell lung cancer (NSCLC) and required for malignant cell survival (Mi et al. 2017). SAGA complex cofactor TRRAP interacts with multiple proteins key to oncogenesis, such as c-Myc and E2F proteins (McMahon et al. 1998). Expression of SAGA *USP22* associates with an oncogenic signature of poor prognosis (Schrecengost et al. 2014). However, recent studies propose a tumor suppressor function of USP22 (Kosinsky et al. 2019), including in Acute Myeloid Leukemia (AML) (Melo-Cardenas et al. 2018). We have previously identified *KAT2A* as a genetic vulnerability in AML, with putative impact on survival and/or differentiation of leukemic cells (Tzelepis et al. 2016). More recently, using a *Kat2a* conditional KO mouse model, we showed that *Kat2a* loss reduces self-renewing AML stem-like cells through impact on transcriptional activity of general metabolic programs, including ribosomal assembly and protein synthesis (Domingues et al. 2020).

Herein, we attempted to characterize the role of KAT2A-containing ATAC and SAGA complexes in normal and leukemic blood cell function. We identified unique ATAC-specific requirements for propagation and differentiation of normal red blood cell progenitors, which did not extend to the SAGA complex. In AML cells, loss of ATAC recapitulated the downregulation of ribosomal protein genes observed upon *Kat2a* loss, establishing a unique metabolic requirement for ATAC that is not mimicked by SAGA. On the other hand, loss of SAGA elements promoted differentiation of AML cells. This opens the possibility of therapeutic targeting of KAT2A activity in the specific context of SAGA. In summary, our work establishes a dichotomy between ATAC and SAGA-centered KAT2A functions that has implications for normal and malignant developmental decisions and can be exploited translationally.

## RESULTS AND DISCUSSION

### KAT2A-containing ATAC and SAGA complexes have unique targets in hematopoietic cells

In order to characterize complex-specific roles of KAT2A in the blood system, we performed chromatin immunoprecipitation followed by next-generation sequencing (ChIP-seq) of ATAC and SAGA-specific subunits (Fig. 1A) in human K562 cells. This is a chronic myelogenous leukemia cell line with multilineage potential (Sutherland et al. 1986): K562 cells can differentiate into red blood cells, platelets or macrophages under defined cytokine conditions; in steady-state, they represent an immature leukemia blast cell with self-renewal properties. K562 cells are thus a good model for understanding mechanisms of normal and malignant blood specification. We made use of specific sera against human ZZZ3 (ATAC-specific) and SPT20 (SAGA-specific) and performed ChIP-seq experiments in duplicate in self-renewing K562 cells. As in previous reports, we found limited overlap between ATAC and SAGA targets (Fig. 1B; Supplemental File 1). Bound peaks were preferentially found in the vicinity of the transcriptional start site (TSS) (Fig. 1C) and were robustly enriched for known ZZZ3 and SPT20 targets, respectively (Fig. S1A-B). Interestingly, ZZZ3 peaks were more proximal than SPT20, contrary to the previously described enhancer association of ATAC complexes in lymphoblast and HeLa cells (Krebs et al. 2011). However, consistent with our results, a distinct ZZZ3 ChIP-seq experiment in human NSCLC cells revealed that ZZZ3 peaks were strongly enriched at regions ±1 kb of TSS (Mi et al. 2017). This suggests that ATAC-complex enhancer region occupancy may be cell type or context dependent. ENCODE experimental enrichment (Kuleshov et al. 2016) of sequence-specific transcription factors within ZZZ3 and SPT20 peaks was clearly distinct (Fig. S1A-B), highlighting complex-specific biologies. These are reflected in distinct gene ontology categories associated with complex-specific peaks, which encompass RNA and ribosomal metabolism in the case of ATAC (Fig. 1D and S1C) and transcriptional activity for SAGA (Fig. S1D). Moreover, SAGA specifically binds red blood cell-associated genes (Fig. 1D), which may indicate a unique role in erythroid differentiation or identity. Critically, we verified the transcriptional significance of unique ATAC and SAGA binding events by confirming loss of target gene expression upon *ZZZ3* or *SUPT20H* knockdown (Fig. 1E), which could be attributed to loss of H3K9ac (Fig. S1E), a well-described target of KAT2A activity (Brownell et al. 1996). ZZZ3 and, to a lesser degree SPT20, were required for propagation of K562 cells (Fig. S1F). Loss of *KAT2A* (Fig. S1G) had an intermediate effect (Fig. S1F), in line with independent activities of each complexes in hematopoietic cell maintenance and/or identity.

### ATAC is selectively required for self-renewal of MOLM13 AML cells

Having established differential molecular requirements for ATAC and SAGA activity in the multipotent K562 line, we explored their contribution to AML biology, where we have established a role for KAT2A (Tzelepis et al. 2016; Domingues et al. 2020). For that, we knocked down *ZZZ3* and *SUPT20H* expression in the MOLM13 AML cell line (Fig. S2A), which we have previously shown to be dependent on KAT2A (Tzelepis et al. 2016). Indeed, loss of *KAT2A* restricts the expansion of MOLM13 cell cultures (Fig. S2B), but in this case its effect seems to reflect at least added activities of ATAC and SAGA complexes (Fig. S2B). Similar to K562 cells, *ZZZ3* knockdown results in downregulation of ribosomal protein genes (Fig. 2A), suggesting a pervasive regulation of biosynthetic activity by the ATAC complex. Similar observations were made in *ZZZ3* and *YEATS2-*depleted lung cancer cell lines (Mi et al. 2017; Mi et al. 2018). High protein synthesis activity is required for leukemia stem cell maintenance (Signer et al. 2014). Significantly, we have observed a reduction in protein synthesis activity in mouse AML cells upon *Kat2a* knockout, which accompanies leukemia stem cell loss (Domingues et al. 2020), and may be attributable to ATAC. SAGA, on the other hand, has no effect on ribosomal genes, but it affects the self-renewal *HOXA* gene signature, which is also down-regulated upon *ZZZ3* knockdown (Fig. 2B). Analysis of the cellular consequences of gene knockdown suggest that ATAC and SAGA act on distinct aspects of leukemia cell maintenance. Loss of SAGA *SUPT20H* induced differentiation of leukemic blasts (Fig. 2C and S2C) and resulted in a trend towards apoptosis (Fig. 2D and S2D), none of which were elicited through loss of *ZZZ3*. In contrast, *ZZZ3* knockdown uniquely affected cell cycle progression, with accumulation of cells in G0/G1, which was not seen with loss of *SUPT20H* (Fig. 2E and S2E), compatible with previous observations in mouse and human cell lines (Orpinell et al. 2010). Also, loss of ATAC-specific YEATS2 results in G1 arrest of lung cancer cells, with no defect in apoptosis (Mi et al. 2017). Overall, the data suggest that while both complexes contribute to leukemia propagation, they affect separate aspects of metabolism and proliferation (ATAC) and cell identity and survival (SAGA), which together explain the KAT2A requirement in leukemia stem cells.

**Figure 2.**
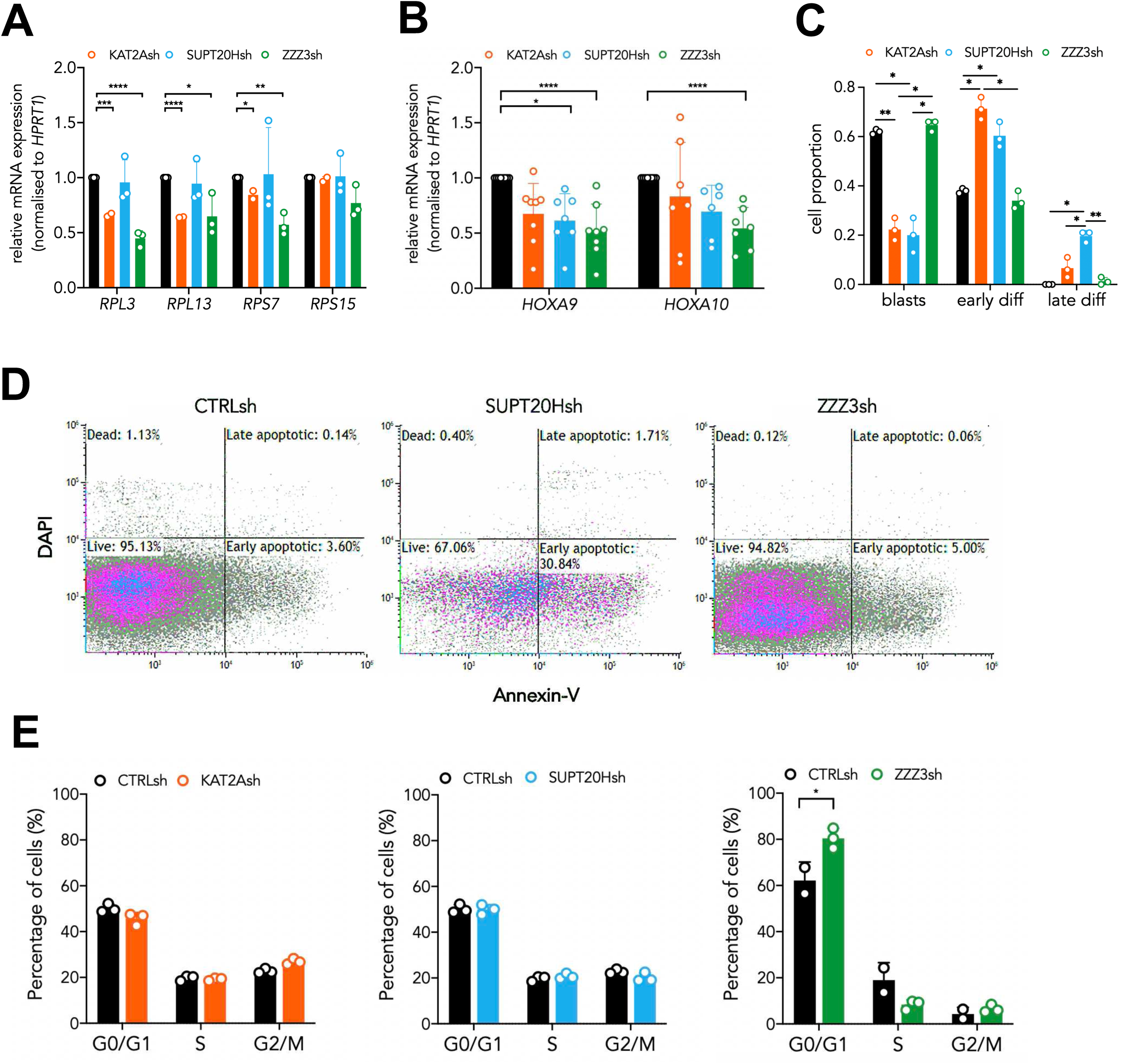
ATAC is selectively required for self-renewal of MOLM13 AML cells. **(A)** Quantitative RT-PCR analysis of ribosomal protein gene expression in MOLM-13 cells transduced with *KAT2Ash*, ZZZ3sh and *SUPT20Hsh*. N ≥3 individual experiments, mean ± SEM of gene expression relative to *CTRLsh*, normalised to *HPRT1* housekeeping gene. Two-tailed t-test for significance *p<0.05, **p<0.01, ***p<0.001. **(B)** Quantitative RT-PCR analysis of self-renewal gene signature in MOLM-13 cells transduced with *KAT2Ash*, ZZZ3sh and *SUPT20Hsh*. N=2 biological replicates, each run as 2 or 3 technical repeats; mean ± SEM of relative gene expression relative to *CTRLsh*, normalised to *HPRT1* housekeeping gene. Two-tailed nested t-test for significance *p<0.05, **p<0.01, ***p<0.001. **(C)** Quantification of blast-like and monocytic differentiated cells in MOLM-13 cultures transduced with *KAT2Ash*, ZZZ3sh and *SUPT20Hsh*. Scoring of 3 randomly selected fields of >100 cells; Nested two-tailed t-test for significance; *p<0.05, **p<0.01, ****p<0.0001. **(D)** Representative flow cytometry plots of apoptosis analysis of MOLM-13 cells transduced with *CTRLsh, SUPT20Hsh* and *ZZZ3sh*. The combination of Annexin V and DAPI staining allows for a distinction between viable cells (double negative), cells in early apoptosis (Annexin V positive), cells in late apoptosis (double positive) and dead cells (DAPI positive). **(E)** Flow cytometry analysis of cell cycle in MOLM-13 cells transduced with *KAT2Ash, ZZZ3sh* and *SUPT20Hsh*. Mean ± SEM of 3 independent experiments. Two-tailed t-test for significance *p<0.05, **p<0.01, ***p<0.001.

### ATAC is selectively required for erythroid specification from cord blood HSC

Given our observations that ATAC and SAGA have distinct effects in leukemic cell fate decisions, we next asked whether the 2 complexes also played different roles in normal hematopoiesis. Of note, KAT2A loss or inactivation has not been associated with hematopoietic defects (Bararia et al. 2016; Domingues et al. 2020). However, the differential averaging *vs* additive contributions of the complexes to global KAT2A activity observed in K562 *vs* MOLM13 cells (Fig. S1F and S2B), leaves open the possibility of ATAC or SAGA being specifically required in normal blood. For this, we transduced normal human CD34+ cord blood (CB) cells with *ZZZ3* or *SUPT20H shRNAs* (Fig. S3A-B) and flow-sorted transduced (GFP+) stem and multipotent progenitor cells (HSC) and lineage-restricted myelo-lymphoid (MLP), megakaryocytic-erythroid (MEP) and granulocytic-monocytic (GMP) progenitors (Fig. 3A). We did not observe differences in the stem and progenitor proportions of transduced cells with either construct (Fig. 3B). In contrast, when assessing cell differentiation and proliferation potential in colony-forming (CFC) progenitor assays (Fig. S3C), we observed a unique defect in erythroid (E) specification from HSC upon *ZZZ3* loss (Fig. 3C), with no changes to generation of mixed-lineage (Mix) or GM colonies. Colony formation from downstream lineage-restricted progenitors was not affected, suggesting a unique requirement in early erythroid commitment. Interestingly, a recent single-cell transcriptomic study detailing the transcriptional trajectory of mouse erythroid cell specification, revealed an association of RNA processing and protein synthesis signatures with early erythroid-committed progenitors (Tusi et al. 2018). This would support a regulatory role for ATAC in the erythroid trajectory through control of protein biosynthetic activity. *SUPT20H* knockdown did not affect colony formation from normal CB stem and progenitor cells (Fig. 3D), suggesting a preferential role in leukemic cells that may be relevant therapeutically. Taken together, these data reveal a unique requirement for ATAC at the level of early human erythroid specification, which led us to revisit the impact of *KAT2A* loss in normal human CB.

**Figure 3.**
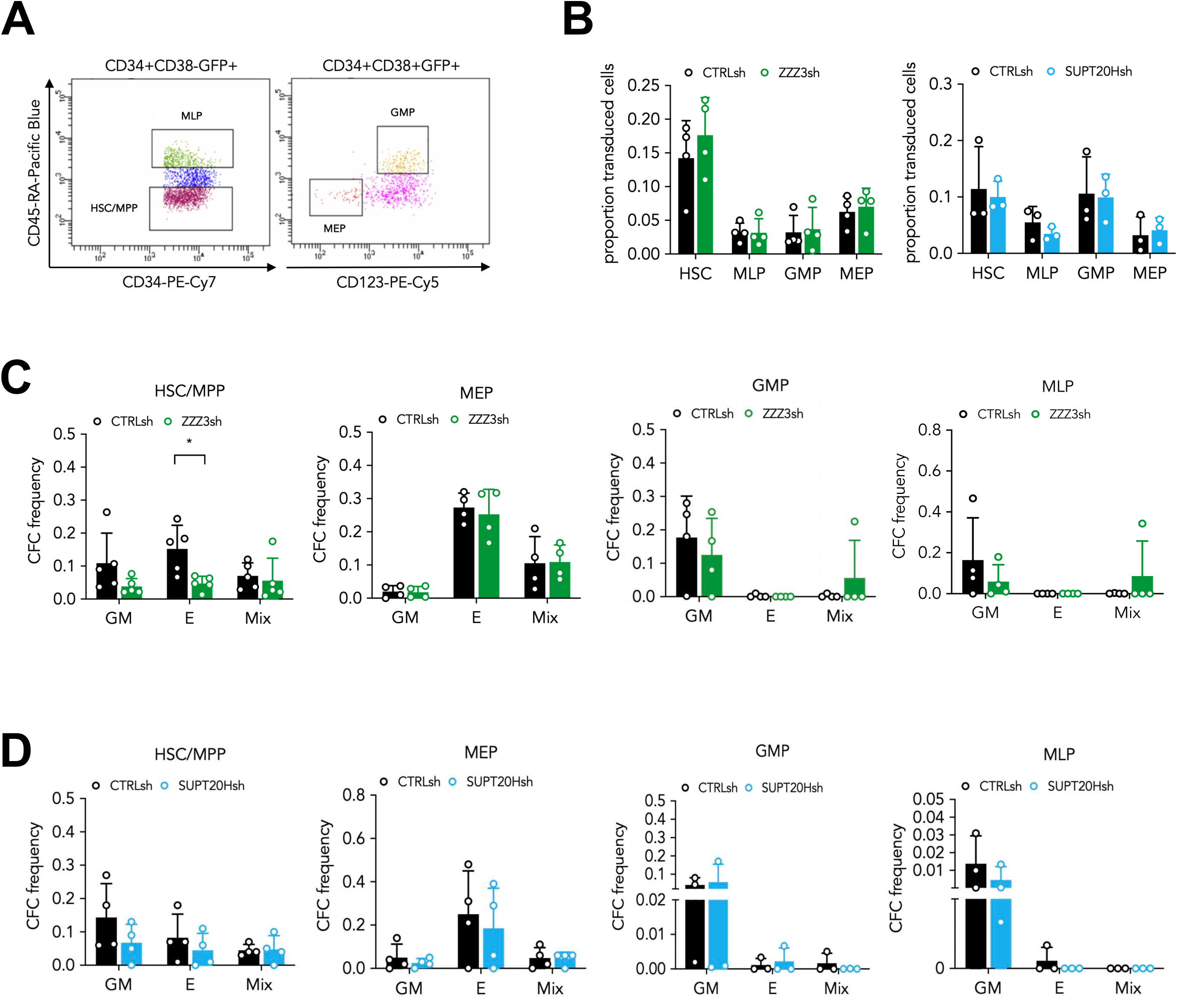
ATAC is selectively required for erythroid specification from CB HSC. **(A)** Representative sorting plot for transduced HSC and progenitor cells from CB. **(B)** Proportion of *ZZZ3sh* (left) and *SUPT20Hsh* (right) transduced CB HSC and progenitors. Mean ± SEM of >3 individual sorting experiments. Two-tailed paired t-test for significance; no significant differences. **(C)** Frequency of colony forming efficiency (CFC) in the HSC/MPP, MEP, GMP and MLP compartments transduced with *ZZZ3sh*. Mean ± SEM of 5 individual CB samples (4 for GMP, MLP). Two-tailed paired t-test for significance; *p<0.05, **p<0.01 **(D)** Frequency of CFC efficiency in the HSC/MPP, MEP, GMP and MLP compartments transduced with *SUPT20Hsh*. Mean ± SEM of 4 individual CB samples. Two-tailed paired t-test for significance; no significant differences.

### KAT2A regulates human CB erythroid progenitor specification and survival

For this, we transduced CD34+ cells with *KAT2A shRNA* (Fig. S4A) and flow-sorted stem and progenitor GFP+ cells, as described (Fig. 3A). We observed a significant decrease in the proportion of KAT2A-depleted MEP (Fig. 4A), suggesting a defect in specification of progenitors committed to the erythroid lineage. Accordingly, we observed a reduction in E colony-formation from HSC (Fig. 4B). Transcriptional analysis of *KAT2A*-knockdown HSC by RNA-sequencing showed an overall reduction in erythroid and megakaryocytic-associated gene signatures (Fig. 4C-D; Supplemental File 2), which is compatible with a defect in specification or survival of MEPs. Indeed, quantification of apoptotic cells revealed a survival defect in both HSC and MEP upon KAT2A depletion (Fig. S4B). There were no changes in cell cycle (Fig. S4C-D). Progenitors affiliated to the granulocyte-monocyte lineage were unaffected in their colony output, survival or proliferation (Fig. 4B and Fig. S4B-C). However, we did observe a mild but significant defect in E colony-formation from MEP (Fig. 4B), which we had not seen upon loss of *ZZZ3* or *SUPT20H* expression. Overall, the results suggest that KAT2A regulates human erythroid biology at multiple levels. Firstly, at the level of specification and/or survival of MEP, which are likely dependent on ATAC activity. Compatible with this view, our inspection of the detailed single-cell profiling of erythroid development by Tusi et al. (2018) (Tusi et al. 2018) noted that both *Kat2a* and *Zzz3*, but no elements of the SAGA complex, were enriched at the transition of multipotent progenitors to the erythroid and megakaryocytic lineages (Fig. S5A; Supplemental File 3). Secondly, KAT2A functions in differentiation of the erythroid lineage, a role that cannot be attributed to ATAC through perturbation of the ZZZ3 subunit, but for which the role of SAGA is ambiguous. Although loss of *SUPT20H* expression did not significantly impair E activity from either HSC or MEP, SPT20 specifically bound erythroid lineage-affiliated *loci* such as *HBB* (Fig. 1D and Fig. S1E), for which expression was down-regulated upon *SUPT20H*, but not *ZZZ3* knockdown, in K562 cells (Fig. 1E). Significantly, erythroid regulator *GATA1* was also mildly down-regulated upon *SUPT20H* knockdown (Fig. S5B), indicating a contribution, albeit likely not an absolute requirement, to regulation of the erythroid differentiation program.

**Figure 4.**
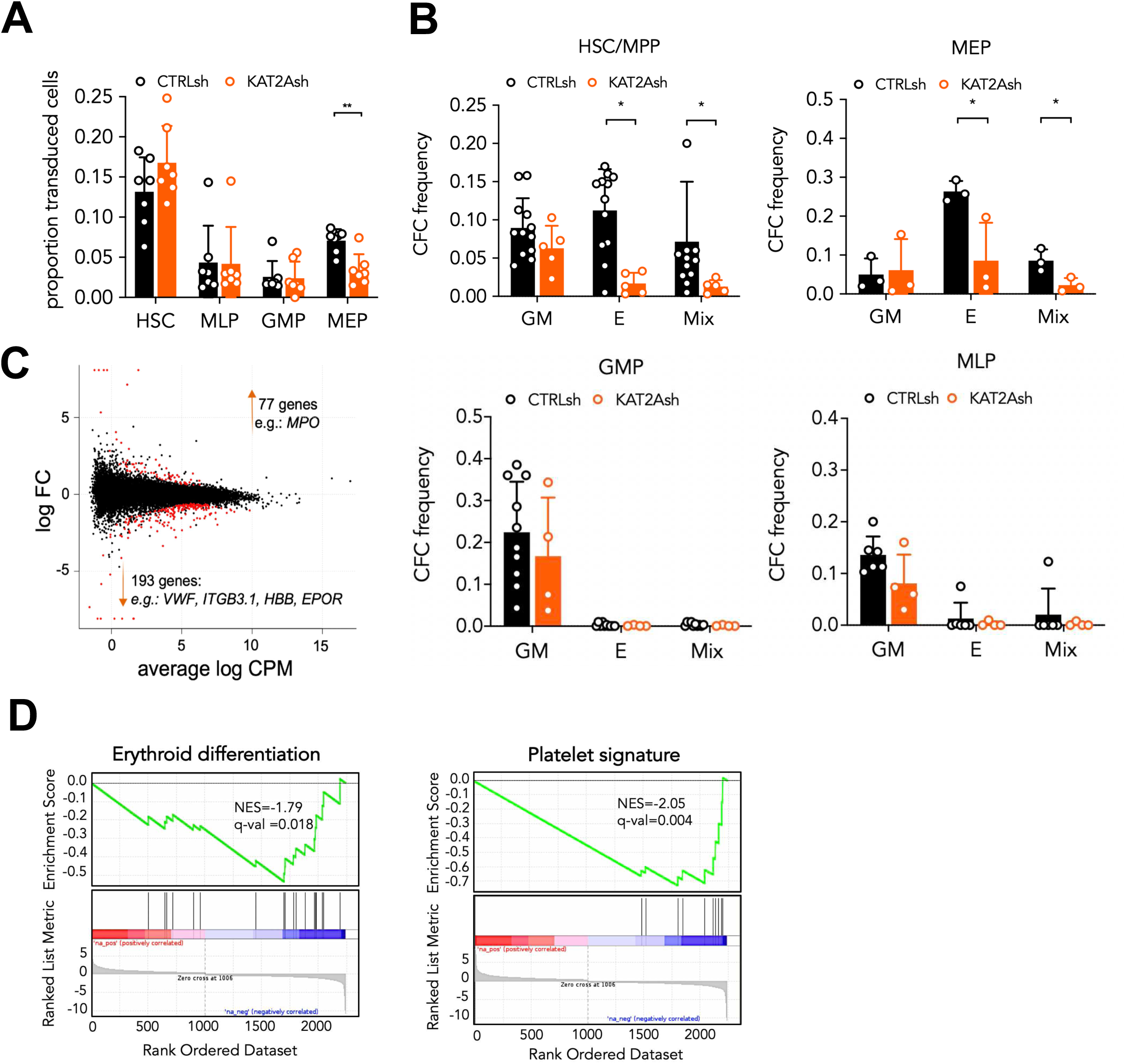
KAT2A regulates human CB erythroid progenitor specification and survival. **(A)** Proportion of *KAT2Ash* transduced CB HSC and progenitors. Mean ± SEM of >7 individual sorting experiments. Two-tailed paired t-test for significance; *p<0.05, **p<0.01. **(B)** Frequency of CFC efficiency in the HSC/MPP, MEP (top), GMP and MLP (bottom) compartments transduced with *KAT2Ash*. Mean ± SEM of 7 individual CB samples. Two-tailed paired t-test for significance; *p<0.05, **p<0.01. **(C)** MA plot of RNA-seq gene expression analysis of DE genes (red) in *CTRLsh vs KAT2A*sh transduced HSC/MPP cells. **(D)** Gene Set Enrichment Analysis (GSEA) plot for erythroid differentiation (right) and platelet signature (left) in the RNA-seq data in (C).

### Loss of SAGA USP22 impacts human erythroid differentiation post-commitment

Inspection of the expression pattern of SAGA-specific elements during *in vitro* maturation of committed erythroid progenitors from human CB (Merryweather-Clarke et al. 2011) shows that several elements associate with late differentiation (Fig. S5C), including members of the Core, but also the SAGA-specific HAT subunit *TADA2B* and the H2B de-ubiquitinase *USP22* (Supplemental File 4). SAGA HAT can exist as a separate complex, at least in *Drosophila* (Soffers et al. 2019), which may explain some level of independence from the Core. On the other hand, the recent elucidation of the structure of yeast SAGA (Papai et al. 2020; Wang et al. 2020) suggests that DUB nucleosome binding is an early event in complex reconfiguration leading to activation of transcription. The structure of the HAT module during the process could not be resolved due to its loose association with the Core, but its activity may be coordinated with the DUB module, as both their dependent histone modifications promote transcription. We thus asked if loss of DUB-*USP22* in human CB could mimic the defects observed in MEP colony-formation upon *KAT2A* knockdown. As before, we transduced human CB CD34+ cells with a lentiviral-delivered targeting shRNA (Fig. 5A) and sorted HSC, MLP, GMP and MEP for downstream analysis. We could not detect differences in the relative representation of stem and progenitor GFP+ cells post-*USP22 shRNA* (Fig. 5B), suggesting that the MEP loss observed upon *KAT2A* knockdown may translate added effects of KAT2A participation in ATAC and SAGA. However, unlike *SUPT20H* knockdown, reduction of *USP22* expression did abrogate E colony formation from MEP (Fig. 5C), critically with no impact on early E colony readout from HSC (Fig. 5C). Colony output from MLP and GMP, and indeed mixed-lineage or GM colony formation from HSC, were not affected (Fig. 5C), indicating a unique contribution of USP22 to expansion and differentiation of MEP. Altogether, the data are compatible with a requirement for SAGA-dependent enzymatic activities on erythroid differentiation post-commitment, which may not be strictly dependent on the association with the Core. This could be because one or both enzymatic modules may exist and interact as independent complexes in at least some cell types (Soffers et al. 2019). Alternatively, the presence of residual SPT20 transcript upon knockdown, rather than knockout, of *SUPT20H* expression, may also allow for sufficient basal SAGA assembly. Nevertheless, a similar level of knockdown triggered differentiation of leukemia cells, indicating differential dependence on normal and malignant hematopoietic cells on SAGA.

**Figure 5.**
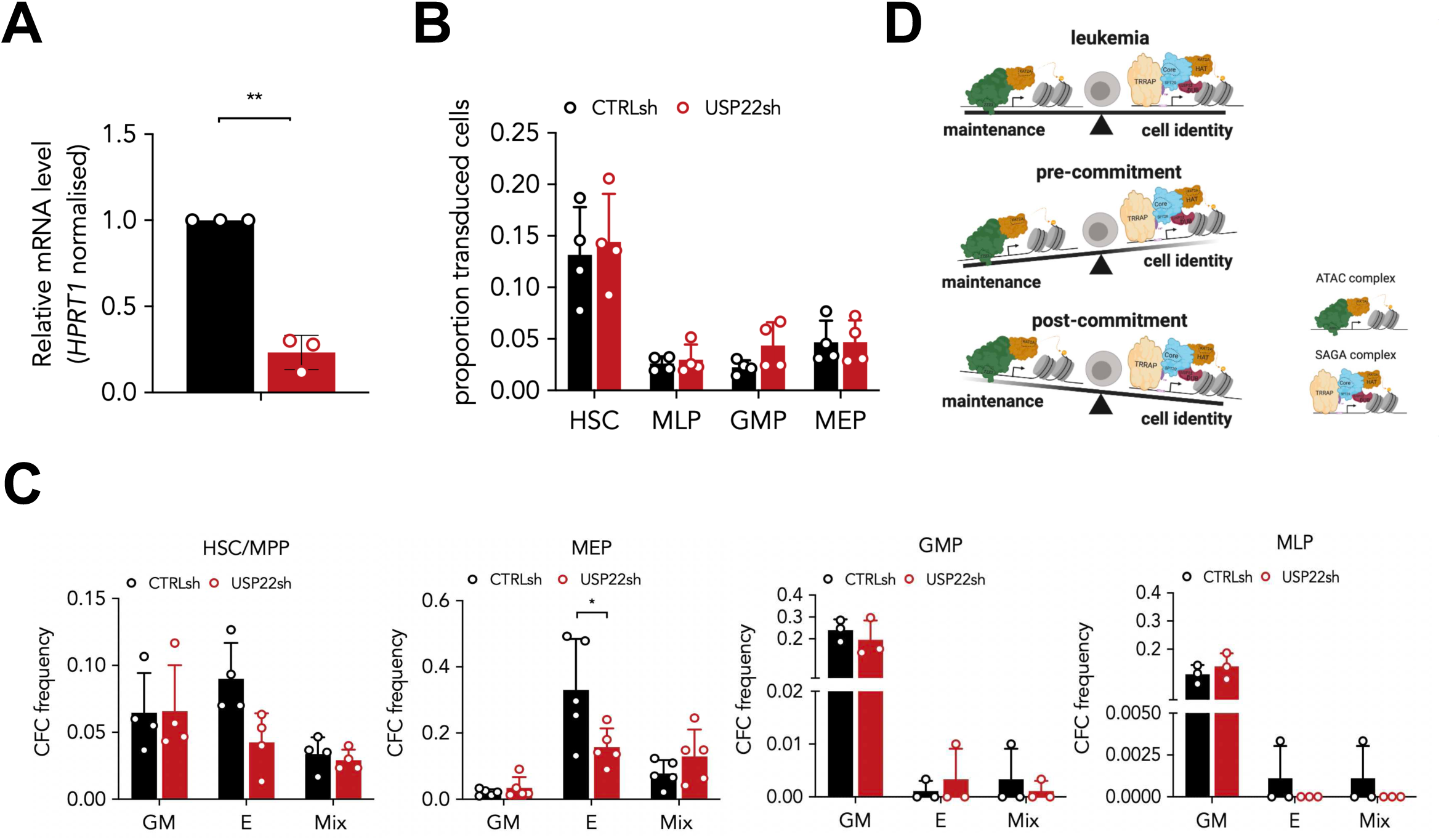
Loss of SAGA does not impact normal human erythroid lineage programmes. **(A)** Quantitative RT-PCR validation of *USP22* knockdown in CB HSCs. Mean ± SEM of 3 individual experiments; gene expression relative to *CTRLsh*, normalised to *HPRT1* housekeeping gene. Paired two-tailed t-test for significance **p<0.01. **(B)** Proportion of *USP22sh* transduced CB HSC and progenitors. Mean ± SEM of >4 individual sorting experiments. Two-tailed paired t-test for significance; no significant differences. **(C)** Frequency of CFC efficiency in the HSC/MPP, MEP (top), GMP and MLP (bottom) compartments transduced with *USP22sh*. Mean ± SEM of 4 individual CB samples. Two-tailed paired t-test for significance; *p<0.05. **(D)** Proposed model for differential dependence of normal and malignant hematopoietic cells on KAT2A-containing ATAC and SAGA complexes.

Altogether, these findings may highlight unique respective roles of ATAC and SAGA in cell maintenance and identity (Fig. 5D). Both aspects are key to leukemia self-perpetuation, justifying the requirement of both complexes in AML. In normal hematopoiesis, maintenance dominates pre-commitment, and identity post-commitment, with ATAC required early and SAGA late in lineage development. The unique requirements of both complexes in the erythroid lineage may reflect the distinctive nature of erythroid/megakaryocytic programs, which separate early from the stem cell root with *de novo* establishment of transcription programs (Notta et al. 2016). Importantly, our data configure targeting of the SAGA Core as a possible anti-leukemia strategy with limited or no effects on normal blood production and good therapeutic tolerance.

## MATERIALS AND METHODS

### Cell lines

K562 and MOLM13 lines (kind gift from Brian Huntly, Cambridge, UK) were maintained in RPMI supplemented with 20% FBS (Fetal Bovine Serum, Invitrogen), 1% Penicillin/Streptomycin/Amphothericin (P/S/A, Invitrogen) and 2mM L-Gln. HEK 293T cells were grown in DMEM supplemented with 10% FBS, and P/S/A and L-Gln as above.

### Human CB CD34+ cells

CB samples were obtained with informed consent under local ethical approval (REC 07-MRE05-44). Mononuclear cells (MNC) isolated by Ficoll-Pacque density gradient (Stemcell Technologies) centrifugation were enriched for CD34^+^ cells using the RosetteSep™ Human CB CD34 Pre-Enrichment Cocktail (Stem Cell Technologies) as per manufacturer’s instructions.

### Lentiviral packaging and transduction

Viral constructs containing control shRNA (CTRLsh, non-eukaryotic gene targeting) or shRNA targeting *KAT2A* (KAT2Ash), *SUPT20H* (SUPT20Hsh) *USP22* (USP22sh), and *ZZZ3* (*ZZZ3sh*) (Supplemental Table 1) were packaged in HEK 293T cells as previously described (Pina et al. 2008), but using Turbofect (Thermo) or trans-IT (Mirus) as lipofection reagents. Cell lines were transduced overnight with 1-2 T75 packaging flask-equivalents (FE)/10^6^ cells and washed the following day as described (Pina et al. 2008). GFP^+^ cells were sorted and used in downstream assays 4 days later. CD34^+^ CB cells were pre-stimulated in serum-free medium (Stemcell Technologies) with SCF, TPO and Flt3L (respectively 200, 20 and 20 ng/mL) for up to 24h and transduced overnight using 2FE/2-3*10^5^ cells. Cells were washed the following day and cultured in half the cytokine concentration for an additional 2-3 days prior to sorting.

### Chromatin immunoprecipitation (ChIP)

For ChIP-sequencing, chromatin was prepared in duplicate from K562 cells (9 sonication cycles, 30”ON/30”OFF) and immunoprecipitated using anti-SPT20 and anti-ZZZ3 sera prepared in the Tora Lab (Krebs et al. 2011). All procedures, including library preparation and sequencing performed as described (Domingues et al. 2020). For ChIP-qPCR, K562 cells were immunoprecipitated with anti-H3K9ac antibody or rabbit IgG (Supplemental Table 2); eluted DNA was diluted and quantified by SYBR green qPCR using 2μl DNA per triplicate reaction (primers in Supplemental Table 3). Peak enrichments relative to rabbit IgG were determined using the -2^ΔΔCt^ method with a reference intergenic region.

### Statistical analysis

Statistical analysis was performed in GraphPad Prism 7 software (GraphPad Software). Data are reported as mean ± SEM. Significance calculated by 2-tailed t-test at p<0.05 (see individual figure legends for details).

### Data deposition

ChIP-seq and RNA-seq data have been deposited in GEO (accession numbers GSE128902 and GSE128512).

## ACKNOWLEDGMENTS

This work was funded by a Rosetrees Trust PhD Studentship to LA (M650) and a Kay Kendall Leukaemia Fund Intermediate Fellowship (KKL888) to CP. Work in the CP lab was also funded by a Leuka John Goldman Fellowship for Future Science (2017) and a Wellcome Trust/University of Cambridge ISSF Grant to CP. SG is funded by a Lady Tata Memorial Trust PhD Studentship, a Trinity Henry Barlow Trust Studentship and by the Cambridge Trust. This study was also supported by NIH RO1 grant, (1R01GM131626-01 to LT), by Agence Nationale de la Recherche (ANR) Programme grants; AAPG2019 PICen to LT, ANR PRCI AAPG2019 EpiCAST to LT, and grant ANR-10-LABX-0030-INRT and a French State fund managed by the ANR under the frame program Investissements d’Avenir ANR-10-IDEX-0002-02 to IGBMC. Samples were provided by the Cambridge Blood and Stem Cell Biobank, which is supported by the Cambridge NIHR BRC Wellcome Trust – MRC Stem Cell Institute and the Cambridge Experimental Cancer Medicine Centre, UK. The authors thank the Cell Phenotyping Hub NIHR BRC for their expert support in flow cytometry and cell sorting.

Contribution: study conception: CP; study design: L.A., C.P.; data collection and assembly: L.A, E.F., S.W., A.F.D., S.G., E.S., L.T.; data analysis: L.A.; E.F.; S.W., R.K., C.P.; data interpretation: L.A., C.P.; manuscript writing: L.A., C.P.; final approval of manuscript: all authors.

## REFERENCES

Bararia D, Kwok HS, Welner RS, Numata A, Sárosi MB, Yang H, Wee S, Tschuri S, Ray D, Weigert O et al. 2016. Acetylation of C/EBPα inhibits its granulopoietic function. Nature communications 7: 10968–10968.

Belluschi S, Calderbank EF, Ciaurro V, Pijuan-Sala B, Santoro A, Mende N, Diamanti E, Sham KYC, Wang X, Lau WWY et al. 2018. Myelo-lymphoid lineage restriction occurs in the human haematopoietic stem cell compartment before lymphoid-primed multipotent progenitors. Nature Communications 9: 4100.

Brownell JE, Zhou J, Ranalli T, Kobayashi R, Edmondson DG, Roth SY, Allis CD. 1996. Tetrahymena histone acetyltransferase A: a homolog to yeast Gcn5p linking histone acetylation to gene activation. Cell 84: 843–851.

Carré C, Ciurciu A, Komonyi O, Jacquier C, Fagegaltier D, Pidoux J, Tricoire H, Tora L, Boros IM, Antoniewski C. 2008. The Drosophila NURF remodelling and the ATAC histone acetylase complexes functionally interact and are required for global chromosome organization. EMBO Rep 9: 187–192.

Carre C, Szymczak D, Pidoux J, Antoniewski C. 2005. The histone H3 acetylase dGcn5 is a key player in Drosophila melanogaster metamorphosis. Mol Cell Biol 25: 8228–8238.

Carrelha J, Meng Y, Kettyle LM, Luis TC, Norfo R, Alcolea V, Boukarabila H, Grasso F, Gambardella A, Grover A et al. 2018. Hierarchically related lineage-restricted fates of multipotent haematopoietic stem cells. Nature 554: 106–111.

Domingues AF, Kulkarni R, Giotopoulos G, Gupta S, Vinnenberg L, Arede L, Foerner E, Khalili M, Adao RR, Johns A et al. 2020. Loss of Kat2a enhances transcriptional noise and depletes acute myeloid leukemia stem-like cells. Elife 9.

Gao B, Kong Q, Zhang Y, Yun C, Dent SYR, Song J, Zhang DD, Wang Y, Li X, Fang D. 2017. The Histone Acetyltransferase Gcn5 Positively Regulates T Cell Activation. J Immunol 198: 3927–3938.

Grant PA, Duggan L, Cote J, Roberts SM, Brownell JE, Candau R, Ohba R, Owen-Hughes T, Allis CD, Winston F et al. 1997. Yeast Gcn5 functions in two multisubunit complexes to acetylate nucleosomal histones: characterization of an Ada complex and the SAGA (Spt/Ada) complex. Genes Dev 11: 1640–1650.

Guelman S, Kozuka K, Mao Y, Pham V, Solloway MJ, Wang J, Wu J, Lill JR, Zha J. 2009. The Double-Histone-Acetyltransferase Complex ATAC Is Essential for Mammalian Development. Molecular and Cellular Biology 29: 1176–1188.

Helmlinger D, Tora L. 2017. Sharing the SAGA. Trends Biochem Sci 42: 850–861.

Jacobsen SEW, Nerlov C. 2019. Haematopoiesis in the era of advanced single-cell technologies. Nat Cell Biol 21: 2–8.

Kikuchi H, Nakayama M, Kuribayashi F, Imajoh-Ohmi S, Nishitoh H, Takami Y, Nakayama T. 2014. GCN5 is essential for IRF-4 gene expression followed by transcriptional activation of Blimp-1 in immature B cells. Journal of Leukocyte Biology 95: 399–404.

Kosinsky RL, Helms M, Zerche M, Wohn L, Dyas A, Prokakis E, Kazerouni ZB, Bedi U, Wegwitz F, Johnsen SA. 2019. USP22-dependent HSP90AB1 expression promotes resistance to HSP90 inhibition in mammary and colorectal cancer. Cell Death & Disease 10: 911.

Koutelou E, Hirsch CL, Dent SY. 2010. Multiple faces of the SAGA complex. Curr Opin Cell Biol 22: 374–382.

Krebs AR, Karmodiya K, Lindahl-Allen M, Struhl K, Tora L. 2011. SAGA and ATAC histone acetyl transferase complexes regulate distinct sets of genes and ATAC defines a class of p300-independent enhancers. Mol Cell 44: 410–423.

Kuleshov MV, Jones MR, Rouillard AD, Fernandez NF, Duan Q, Wang Z, Koplev S, Jenkins SL, Jagodnik KM, Lachmann A et al. 2016. Enrichr: a comprehensive gene set enrichment analysis web server 2016 update. Nucleic Acids Res 44: W90–97.

Liu G, Zheng X, Guan H, Cao Y, Qu H, Kang J, Ren X, Lei J, Dong M-Q, Li X et al. 2019. Architecture of Saccharomyces cerevisiae SAGA complex. Cell Discovery 5: 25.

Martinez-Cerdeno V, Lemen JM, Chan V, Wey A, Lin W, Dent SR, Knoepfler PS. 2012. N-Myc and GCN5 regulate significantly overlapping transcriptional programs in neural stem cells. PLoS One 7: e39456.

McMahon SB, Van Buskirk HA, Dugan KA, Copeland TD, Cole MD. 1998. The novel ATM-related protein TRRAP is an essential cofactor for the c-Myc and E2F oncoproteins. Cell 94: 363–374.

Melo-Cardenas J, Xu Y, Wei J, Tan C, Kong S, Gao B, Montauti E, Kirsammer G, Licht JD, Yu J et al. 2018. USP22 deficiency leads to myeloid leukemia upon oncogenic Kras activation through a PU.1-dependent mechanism. Blood 132: 423–434.

Merryweather-Clarke AT, Atzberger A, Soneji S, Gray N, Clark K, Waugh C, McGowan SJ, Taylor S, Nandi AK, Wood WG et al. 2011. Global gene expression analysis of human erythroid progenitors. Blood 117: e96–108.

Mi W, Guan H, Lyu J, Zhao D, Xi Y, Jiang S, Andrews FH, Wang X, Gagea M, Wen H et al. 2017. YEATS2 links histone acetylation to tumorigenesis of non-small cell lung cancer. Nature Communications 8: 1088.

Mi W, Zhang Y, Lyu J, Wang X, Tong Q, Peng D, Xue Y, Tencer AH, Wen H, Li W et al. 2018. The ZZ-type zinc finger of ZZZ3 modulates the ATAC complex-mediated histone acetylation and gene activation. Nat Commun 9: 3759.

Moris N, Edri S, Seyres D, Kulkarni R, Domingues AF, Balayo T, Frontini M, Pina C. 2018. Histone Acetyltransferase KAT2A Stabilizes Pluripotency with Control of Transcriptional Heterogeneity. STEM CELLS 36: 1828–1838.

Notta F, Zandi S, Takayama N, Dobson S, Gan OI, Wilson G, Kaufmann KB, McLeod J, Laurenti E, Dunant CF et al. 2016. Distinct routes of lineage development reshape the human blood hierarchy across ontogeny. Science 351: aab2116.

Orpinell M, Fournier M, Riss A, Nagy Z, Krebs AR, Frontini M, Tora L. 2010. The ATAC acetyl transferase complex controls mitotic progression by targeting non-histone substrates. The EMBO journal 29: 2381–2394.

Papai G, Frechard A, Kolesnikova O, Crucifix C, Schultz P, Ben-Shem A. 2020. Structure of SAGA and mechanism of TBP deposition on gene promoters. Nature 577: 711–716.

Pina C, May G, Soneji S, Hong D, Enver T. 2008. MLLT3 regulates early human erythroid and megakaryocytic cell fate. Cell stem cell 2: 264–273.

Riss A, Scheer E, Joint M, Trowitzsch S, Berger I, Tora L. 2015. Subunits of ADA-two-A-containing (ATAC) or Spt-Ada-Gcn5-acetyltrasferase (SAGA) Coactivator Complexes Enhance the Acetyltransferase Activity of GCN5. J Biol Chem 290: 28997–29009.

Schrecengost RS, Dean JL, Goodwin JF, Schiewer MJ, Urban MW, Stanek TJ, Sussman RT, Hicks JL, Birbe RC, Draganova-Tacheva RA et al. 2014. USP22 regulates oncogenic signaling pathways to drive lethal cancer progression. Cancer Res 74: 272–286.

Signer RA, Magee JA, Salic A, Morrison SJ. 2014. Haematopoietic stem cells require a highly regulated protein synthesis rate. Nature 509: 49–54.

Soffers JHM, Li X, Saraf A, Seidel CW, Florens L, Washburn MP, Abmayr SM, Workman JL. 2019. Characterization of a metazoan ADA acetyltransferase complex. Nucleic Acids Res 47: 3383–3394.

Spedale G, Timmers HTM, Pijnappel WWMP. 2012. ATAC-king the complexity of SAGA during evolution. Genes Dev 26: 527–541.

Suganuma T, Gutierrez JL, Li B, Florens L, Swanson SK, Washburn MP, Abmayr SM, Workman JL. 2008. ATAC is a double histone acetyltransferase complex that stimulates nucleosome sliding. Nat Struct Mol Biol 15: 364–372.

Sun X-J, Man N, Tan Y, Nimer SD, Wang L. 2015. The Role of Histone Acetyltransferases in Normal and Malignant Hematopoiesis. In Front Oncol, p. 108.

Sutherland JA, Turner AR, Mannoni P, McGann LE, Turc JM. 1986. Differentiation of K562 leukemia cells along erythroid, macrophage, and megakaryocyte lineages. J Biol Response Mod 5: 250–262.

Tusi BK, Wolock SL, Weinreb C, Hwang Y, Hidalgo D, Zilionis R, Waisman A, Huh JR, Klein AM, Socolovsky M. 2018. Population snapshots predict early haematopoietic and erythroid hierarchies. Nature 555: 54–60.

Tzelepis K, Koike-Yusa H, De Braekeleer E, Li Y, Metzakopian E, Dovey OM, Mupo A, Grinkevich V, Li M, Mazan M et al. 2016. A CRISPR Dropout Screen Identifies Genetic Vulnerabilities and Therapeutic Targets in Acute Myeloid Leukemia. Cell Rep 17: 1193–1205.

Wang H, Dienemann C, Stützer A, Urlaub H, Cheung ACM, Cramer P. 2020. Structure of the transcription coactivator SAGA. Nature 577: 717–720.

Wang Y, Guo YR, Liu K, Yin Z, Liu R, Xia Y, Tan L, Yang P, Lee JH, Li XJ et al. 2017a. KAT2A coupled with the alpha-KGDH complex acts as a histone H3 succinyltransferase. Nature 552: 273–277.

Wang Y, Yun C, Gao B, Xu Y, Zhang Y, Wang Y, Kong Q, Zhao F, Wang CR, Dent SYR et al. 2017b. The Lysine Acetyltransferase GCN5 Is Required for iNKT Cell Development through EGR2 Acetylation. Cell Rep 20: 600–612.

Xu W, Edmondson DG, Evrard YA, Wakamiya M, Behringer RR, Roth SY. 2000. Loss of Gcn5l2 leads to increased apoptosis and mesodermal defects during mouse development. Nature Genetics 26: 229–232.

