## Supplemental methods, tables, figures and legends for "Unique roles of ATAC and SAGA - KAT2A complexes in normal and malignant hematopoiesis"

### SUPPLEMENTAL MATERIAL
### for

#### *Unique roles of ATAC and SAGA - KAT2A complexes in normal and malignant hematopoiesis*

Liliana Arede[1,2], Elena Foerner[1*], Selinde Wind[1*], Rashmi Kulkarni[1], Ana Filipa Domingues[1,7], Svenja Kleinwaechter[1], Shikha Gupta[1,2], Elisabeth Scheer[3,4,5], Laszlo Tora[3,4,5], Cristina Pina[2,6,8]

[1]Department of Haematology and [2]Department of Genetics, University of Cambridge, Cambridge, United Kingdom; [3]Institut de Génétique et de Biologie Moléculaire et Cellulaire, 67404 Illkirch, France; [4]Centre National de la Recherche Scientifique (CNRS), UMR7104, 67404 Illkirch, France; [5]Institut National de la Santé et de la Recherche Médicale (INSERM), U1258, 67404 Illkirch, France; [6]College of Health and Life Sciences, Division of Biosciences, Brunel University London, Uxbridge UB8 3PH

[7]Present address: Wellcome Trust Cambridge Stem Cell Institute and Department of Haematology, University of Cambridge, Cambridge, United Kingdom *Contributed equally

KEYWORDS: ATAC; SAGA; Self-renewal; Differentiation; Leukemia; Normal haematopoiesis

[8]Correspondence to:

Cristina Pina, D.Phil.
College of Health and Life Sciences, Division of Biosciences
Brunel University London
Uxbridge UB8 3PH
London
United Kingdom
+44 (0)1895 266676

#### SUPPLEMENTARY MATERIALS AND METHODS

##### Flow cytometry

CD34+ CB cells or cell lines were stained as described (Pina et al. 2008): cell surface antibodies are listed in Supplemental Table 4; quantification of apoptosis used Annexin V (Thermo); cell cycle profiles were obtained using Hoechst 33342 (Thermo). Flow analysis data was acquired on Gallios (Beckman Coulter) and Attune (Thermo) instruments and analysed in Kaluza (Beckman Coulter). Cells were sorted on a FACSAria™ Fusion or Influx instruments (BD).

##### Colony-forming cell (CFC) assays

CFC assays of CB stem and progenitor cells were performed in StemMACS-HSC-CFU complete medium (Miltenyi Biotech) and scored 10–12 days after plating.

##### Cytospins

Cells were centrifuged onto slides for 5min at 700rpm, stained with rapid Romanowsky stain pack as per manufacturer's instructions and fixed with Depex mounting medium.

##### Quantitative Real time PCR (Q-RT-PCR)

RNA extraction, cDNA synthesis and Q-PCR analysis were performed as described (Domingues et al. 2020). Primers and Taqman probes assays (Thermo) are listed in Supplemental Tables 5 and 6, respectively. Relative gene expression calculated by the $2^{-\Delta\Delta Ct}$ method using *HPRT1* as reference.

##### RNA sequencing

RNA was extracted from *CTRLsh* or *KAT2Ash* HSCs obtained from 2 individual donors using Trizol reagent (Thermo) and linear polyacrylamide (Sigma) as a carrier. RNA-seq libraries were prepared at the Cambridge Stem Cell Institute Genomics Core Facility using the Ovation RNA-seq kit (NuGen) with incorporated DNase treatment, as per manufacturer's instructions. Libraries were sequenced on an Illumina HiSeq4000 instrument at the CRUK Cambridge Research Institute Genomics Core Facility using 50bp single-end reads. The raw fastq files were processed as per the RSEM v1.2.31 workflow and aligned to the reference human genome assembly

GRCh37. Differentially expressed genes were obtained at 10% FDR using the R packge edgeR. Gene signatures of down-regulated genes in *KAT2Ash* HSC for gene set enrichment analysis were obtained from MSigDB (Subramanian et al. 2005). Erythroid differentiation (NES= -1.79, q-val=0.018) and platelet biology (NES= -2.05, q-val=0.004).

#### SUPPLEMENTAL TABLES

***Supplemental Table 1.*** *Sequences of shRNA constructs.*

| Oligonucleotide | Sequence (5' to 3') |
|---|---|
| CTRLsh | TCAACAAGATGAAGAGCACCAAGGGATCCGTTGGTGCTCTTCATCTT GTTGTTTTTC |
| KAT2Ash | TGCTGAACTTTGTGCAGTACAAGGGATCCGTTGTACTGCACAAAGTT CAGCTTTTTC |
| SUPT20Hsh | TCCATCAAGTATTCCTCGGAAAGGGATCCTTTCCGAGGAATACTTGA TGGTTTTTT |
| USP22sh | TAGCTACCAGGAGTCCACAAAGGGGATCCCTTTGTGGACTCCTGGT AGCTTTTTTTC |
| ZZZ3sh | TGCATCAGATGACGAAAGTATTGGGATCCAATACTTTCGTCATCTGA TGCTTTTTTC |

***Supplemental Table 2.*** *Antibodies used in chromatin immunoprecipitation (ChIP).*

| Antibody | Catalogue # | Supplier |
|---|---|---|
| H3K9ac | 07-352 | Milipore |
| Rabbit IgG | 12-370 | Milipore |

***Supplemental Table 3.*** *Sequences of ChIP-qPCR primers.*

| Gene | Forward | Reverse |
|---|---|---|
| *HBB* | *GCCATCCATTTTTCTTAATTCTGAG* | *TGAGGGCACCATTAGCCAG* |
| *Intergenic region* | *TGGTTTGGAGTGGGTGCT* | *TCCTGCTCTCCGTCACCT* |
| *RPS7* | *CCTGCTCTCCGACAGAACTT* | *CGGGTAATCGGCTGTATCCC* |

***Supplemental Table 4.*** *Antibodies used in flow cytometry analysis and cell sorting.*

| Antibody | Fluorochrome | Catalogue # | Clone | Dilution | Supplier |
|---|---|---|---|---|---|
| CD34 | PE-Cy7 | 343515 | 581 | 1:200 | BioLegend |
| CD38 | PE | 356603 | HB-7 | 1:100 | BioLegend |

| | | | | | |
|---|---|---|---|---|---|
| CD45RA | APC | 304118 | HI100 | 1:100 | BioLegend |
| CD123 | PE-Cy5 | 306008 | HI264 | 1:100 | BioLegend |
| Hoechst 33352 | - | H3570 | - | | LifeTechnologies |
| Annexin V | APC | 640941 | - | 1:100 | BioLegend |

***Supplemental Table 5.*** *Sequences of qRT-PCR primers.*

| Gene | Forward | Reverse |
|---|---|---|
| *HBB* | *AGGAGAAGTCTGCCGTTACTG* | *CCGAGCACTTTCTTGCCATGA* |
| *HOXA10* | *GAGAGCAGCAAAGCCTCGC* | *CCAGTGTCTGGTGCTTCGTG* |
| *HOXA9* | *GGTGACTGTCCCACGCTTGAC* | *GAGTGGAGCGCGCATGAAG* |
| *HPRT1* | *CCTGGCGTCGTGATTAGTGAT* | *TCGAGCAAGACGTTCAGTCC* |
| *KAT2A* | *CCCGCTACGAAACCACTCAT* | *GCATGGACAGGAATTTGGGGA* |
| *RPL13* | *CGCAGGAGCCGCAGG* | *CTGCCAGTCCTTGTGGAAGT* |
| *RPS7* | *CCCAGGAGCCGTACTCTGA* | *GCCATCTAGTTTGACGCGGA* |
| *SUPT20H* | *CCCTTAAATCTACTCCAGCTTCCAG* | *TTGACTGGTTGAACCTTGCTC* |
| *USP22* | *GAGGCCATGGACGCCG* | *AGATACAGGACTTGGCCTTGC* |
| *ZZZ3* | GGACAGCAAAACAGGTTGCC | GTGCTGTCGTCTGCTTGTTG |

***Supplemental Table 6.*** *Taqman probes.*

| Gene | Catalogue# | Supplier |
|---|---|---|
| *GATA1* | *Hs01085823_m1* | Thermo |
| *HPRT1* | *Hs02800695_m1* | Thermo |
| *KAT2A* | *Hs00221499_m1* | Thermo |
| *RPL3* | *Hs01581771_g1* | Thermo |
| *RPL15* | *Hs04334752_g1* | Thermo |

#### SUPPLEMENTAL FILES

**Supplemental File 1:** ZZZ3 and SPT20 ChIP-seq peaks in human K562 cells

**Supplemental File 2:** RNA-seq differentially expressed genes in CTRLsh *vs* KAT2Ash HSCs (10% FDR)

**Supplemental File 3:** Enriched genes in erythroid-basophil-megakaryocyte-biased progenitors (EBMP) as per detailed single-cell profiling of erythroid development by Tusi et al. (2018). EBMP region selected using SPRING tool (https://kleintools.hms.harvard.edu/paper_websites/tusi_et_al/).

**Supplemental File 4:** Genes enriched in Intermediate Erythroblasts (IntE) as defined in a microarray study of human erythroid differentiation by (Merryweather-Clarke et al. 2011). Cluster 17 obtained from the Human Erythroblast Maturation database (https://cellline.molbiol.ox.ac.uk/eryth/index.html).

#### REFERENCES


Domingues AF, Kulkarni R, Giotopoulos G, Gupta S, Vinnenberg L, Arede L, Foerner E, Khalili M, Adao RR, Johns A et al. 2020. Loss of Kat2a enhances transcriptional noise and depletes acute myeloid leukemia stem-like cells. *Elife* **9**.

Kuleshov MV, Jones MR, Rouillard AD, Fernandez NF, Duan Q, Wang Z, Koplev S, Jenkins SL, Jagodnik KM, Lachmann A et al. 2016. Enrichr: a comprehensive gene set enrichment analysis web server 2016 update. *Nucleic Acids Res* **44**: W90-97.

Merryweather-Clarke AT, Atzberger A, Soneji S, Gray N, Clark K, Waugh C, McGowan SJ, Taylor S, Nandi AK, Wood WG et al. 2011. Global gene expression analysis of human erythroid progenitors. *Blood* **117**: e96-108.

Pina C, May G, Soneji S, Hong D, Enver T. 2008. MLLT3 regulates early human erythroid and megakaryocytic cell fate. *Cell stem cell* **2**: 264-273.

Subramanian A, Tamayo P, Mootha VK, Mukherjee S, Ebert BL, Gillette MA, Paulovich A, Pomeroy SL, Golub TR, Lander ES et al. 2005. Gene set enrichment analysis: A knowledge-based approach for interpreting genome-wide expression profiles. *Proceedings of the National Academy of Sciences* **102**: 15545-15550.

Tusi BK, Wolock SL, Weinreb C, Hwang Y, Hidalgo D, Zilionis R, Waisman A, Huh JR, Klein AM, Socolovsky M. 2018. Population snapshots predict early haematopoietic and erythroid hierarchies. *Nature* **555**: 54-60.

#### SUPPLEMENTAL FIGURES

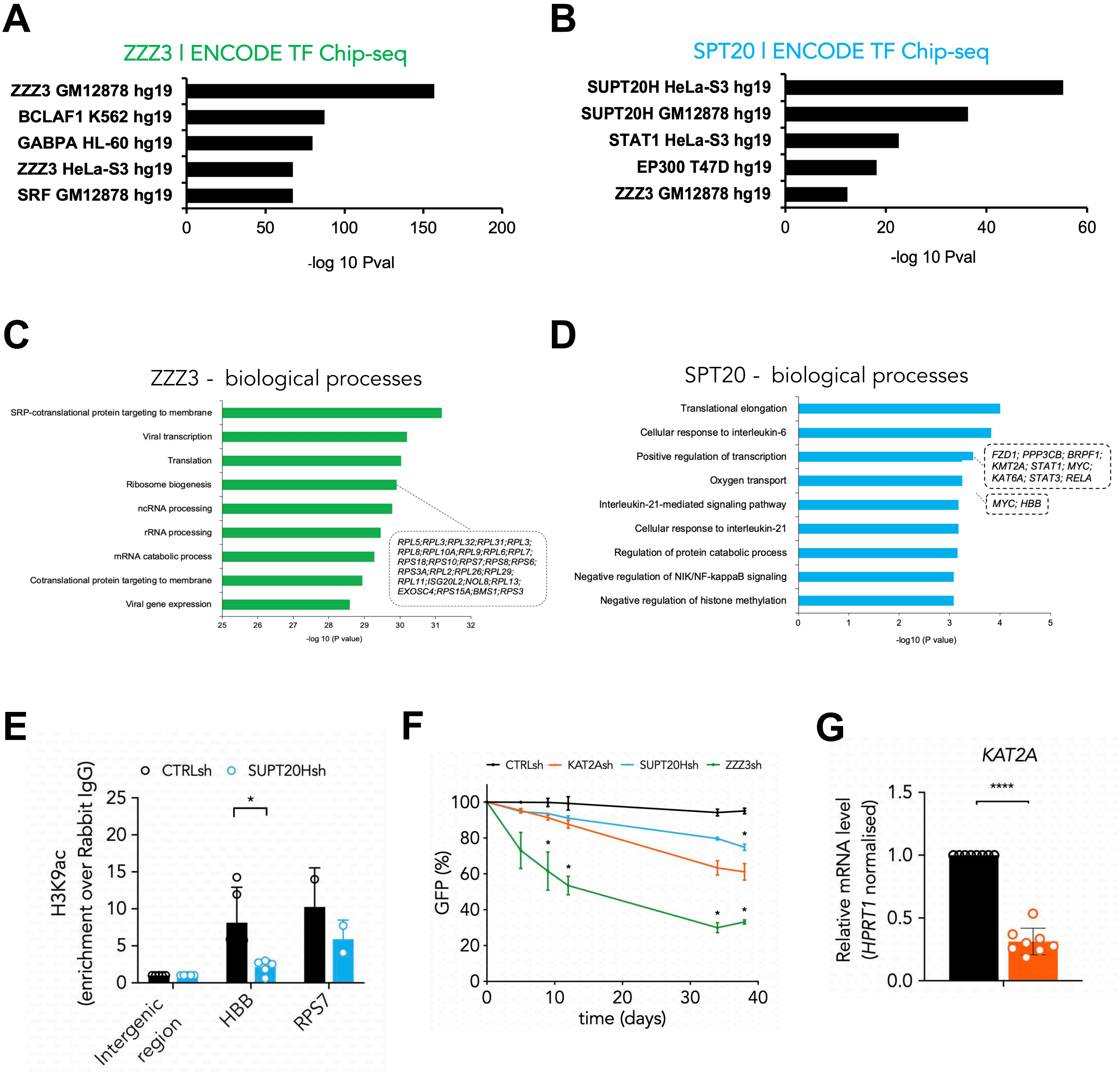


**Figure S1. KAT2A-containing ATAC and SAGA complexes have unique functional associations in K562 cells. (relates to Figure 1)**

**(A-B)** Specificity of ATAC (ZZZ3) and SAGA (SPT20) ChIP-seq targets against ENCODE; data as retrieved by EnrichR online annotation tool (Kuleshov et al., 2016).

**(C-D)** Top Gene Ontology (GO) associations of ZZZ3 and SPT20 ChIP-seq targets on biological processes as calculated by EnrichR (Kuleshov et al., 2016).

**(E)** H3K9ac ChIP-qPCR analysis of SPT20 targets upon knockdown in K562 cells. N≥2 independent experiments. Mean ± SEM of enrichment relative to rabbit-IgG, with normalisation to intergenic region. Two-tailed t-test for significance *p<0.05.

**(F)** Growth curve of K562 cells transduced with shRNA constructs against *ZZZ3*, *SUPT20H* and *KAT2A*. Mean ± SEM of 3 independent experiments. ANOVA for mixed effects analysis significance *p<0.05, **p<0.01, ***p<0.001.

**(G)** Quantitative RT-PCR validation of *KAT2A* knockdown in K562 cells. Mean ± SEM of 8 individual experiments; gene expression relative to CTRLsh, normalised to *HPRT1* housekeeping gene. Paired two-tailed t-test for significance ***p<0.001.

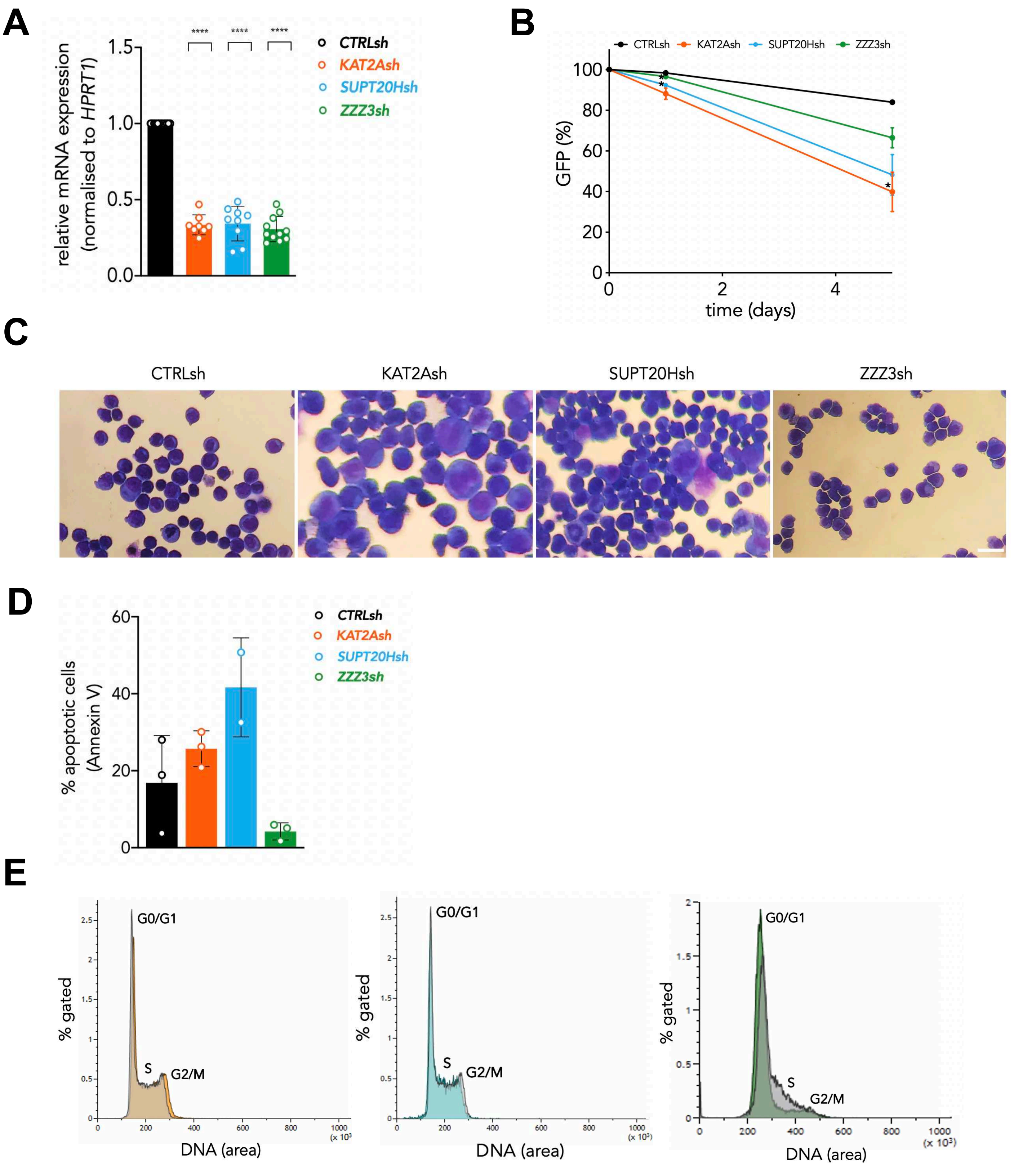


**Figure S2. Loss of SAGA subunits increase differentiation of MOLM13 AML cells. (relates to Figure 2)**

**(A)** Quantitative RT-PCR validation of KAT2A, ZZZ3 and SUPT20H knockdown in MOLM-13 cells. Mean ± SEM of ≥3 individual experiments; gene expression relative to CTRLsh, normalised to *HPRT1* housekeeping gene. Two-tailed t-test for significance *p<0.05, **p<0.01, ***p<0.001.

**(B)** Growth curve of MOLM-13 cells transduced with shRNA constructs against KAT2A, ZZZ3 and SUPT20H. Mean ± SEM of 3 independent experiments. is shown. ANOVA for mixed effects analysis significance *p<0.05, **p<0.01).

**(C)** Representative photographs of MOLM-13 cytospins.

**(D)** Flow cytometry analysis of apoptosis in MOLM-13 cells transduced with shRNA constructs against KAT2A, ZZZ3 and SUPT20H. Mean ± SEM of % Annexin V positive cells in 3 independent experiments. Two-tailed t-test for significance; no significant changes.

**(E)** Representative flow cytometry overlays of cell cycle analysis in MOLM-13 cells transduced with shRNA constructs against KAT2A, ZZZ3 and SUPT20H.

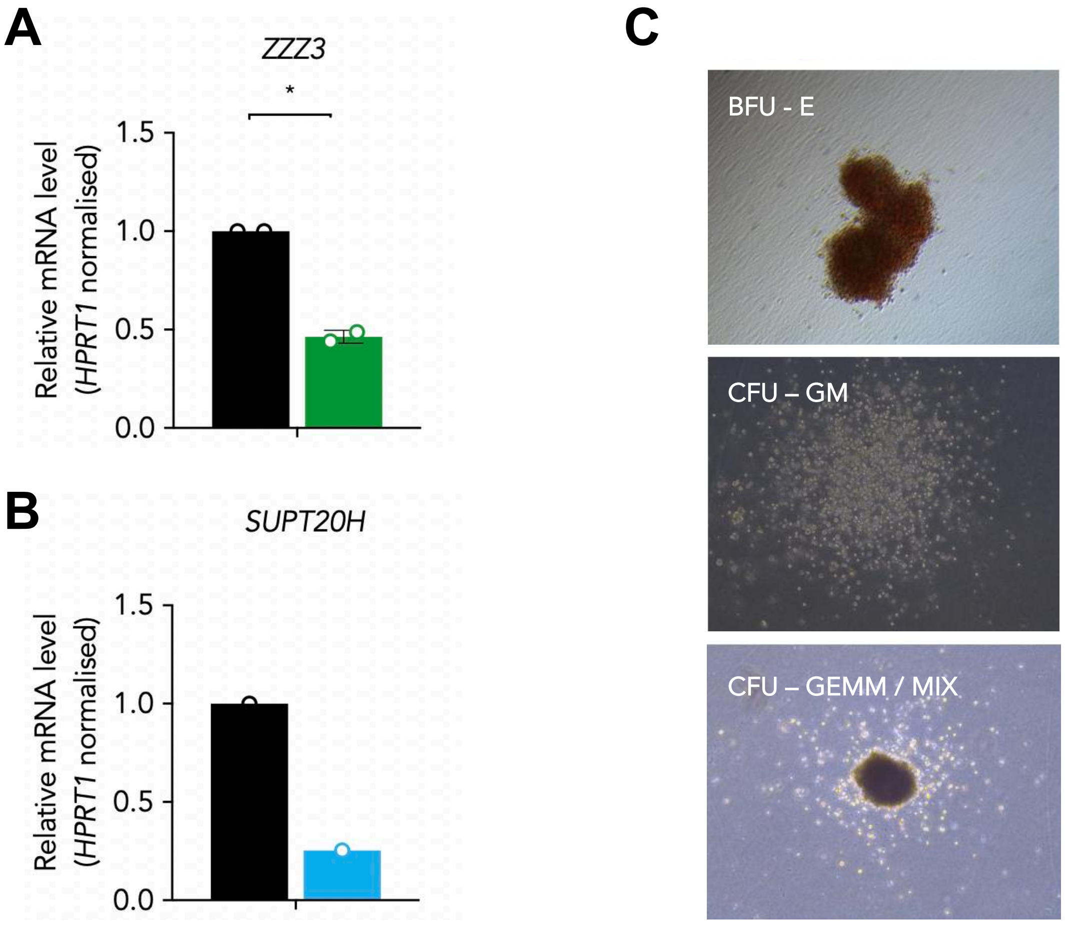


**Figure S3. Investigation of ZZZ3 and SUPT20H knockdown in human cord blood. (relates to Figure 3)**
**(A)** Quantitative RT-PCR validation of *ZZZ3* knockdown in human cord blood HSC. Mean ± SEM of 2 individual experiments; gene expression relative to CTRLsh, normalised to *HPRT1* housekeeping gene. Paired two-tailed t-test for significance $^*p<0.05$.
**(B)** Quantitative RT-PCR validation of *SUPT20H* knockdown in human cord blood HSC. Representative experiment.
**(C)** Photographs of representative colony morphologies from transduced human cord blood HSC and progenitors. BFU-E: Boost-forming unit – erythrocyte colony. CFU-GM: Colony-forming unit - granulocyte monocyte colony; CFU-GEMM or mixed: Colony-forming unit – granulocyte erythrocyte monocyte megakaryocyte colony.

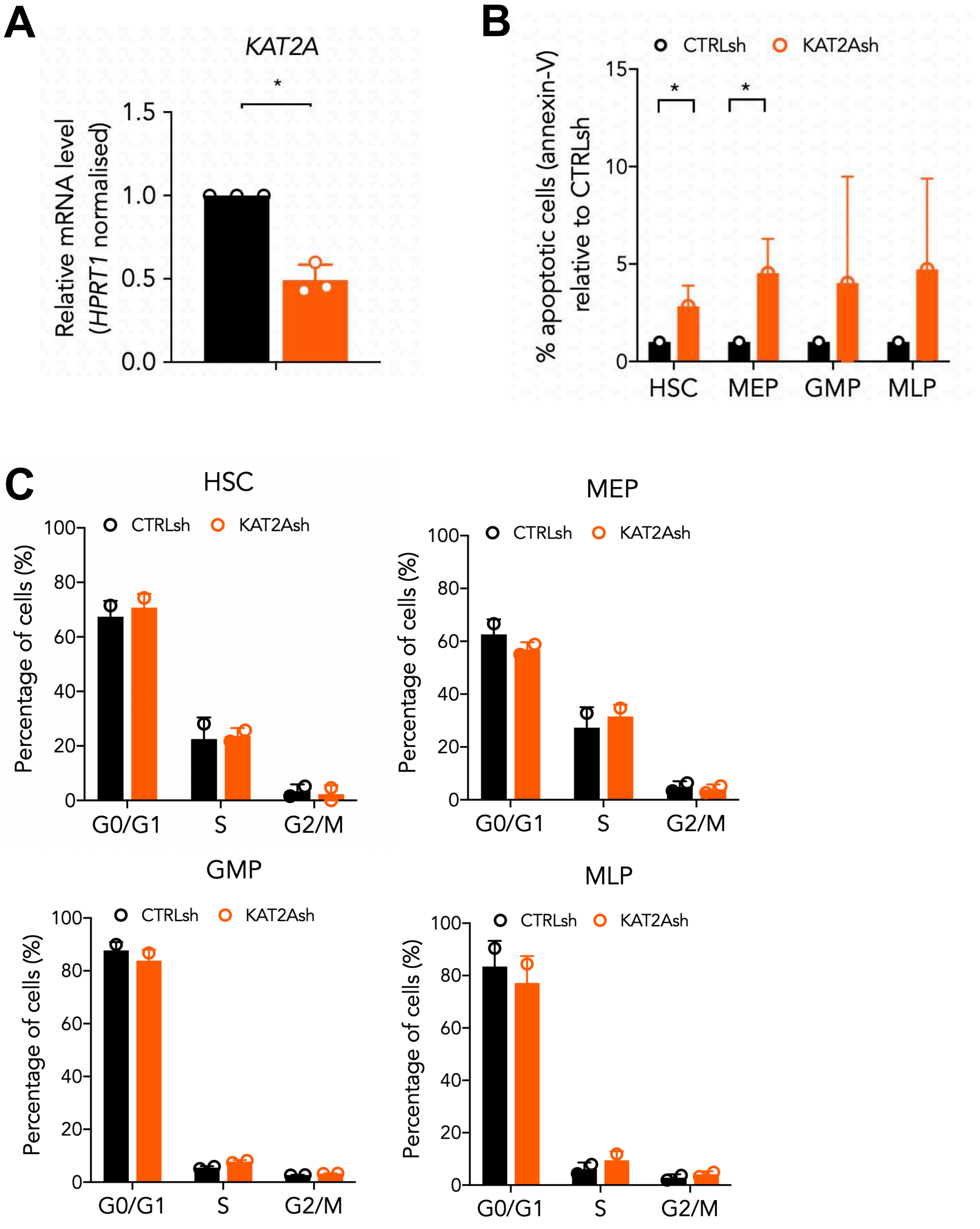


**Figure S4. KAT2A is required for human cord blood progenitor survival and erythroid specification with no impact on lymphoid and myeloid potential. (relates to Figure 4)**
**(A)** Quantitative RT-PCR validation of *KAT2A* knockdown in human cord blood HSC. Mean ± SEM of 3 individual experiments; gene expression relative to CTRLsh, normalised to *HPRT1* housekeeping gene. Paired two-tailed t-test for significance *$p<0.05$.
**(B)** Flow cytometry analysis of apoptosis by Annexin-V staining. N=3 individual cord blood samples; mean ± SEM. Two-tailed paired t-test for significance; *$p<0.05$.
**(C)** Flow cytometry analysis of cell cycle in HSC/MPP, MEP, GMP and MLP transduced cells with KAT2Ash vs CTRLsh. N=3 individual cord blood samples; mean ± SEM. Two-tailed paired t-test not significant.

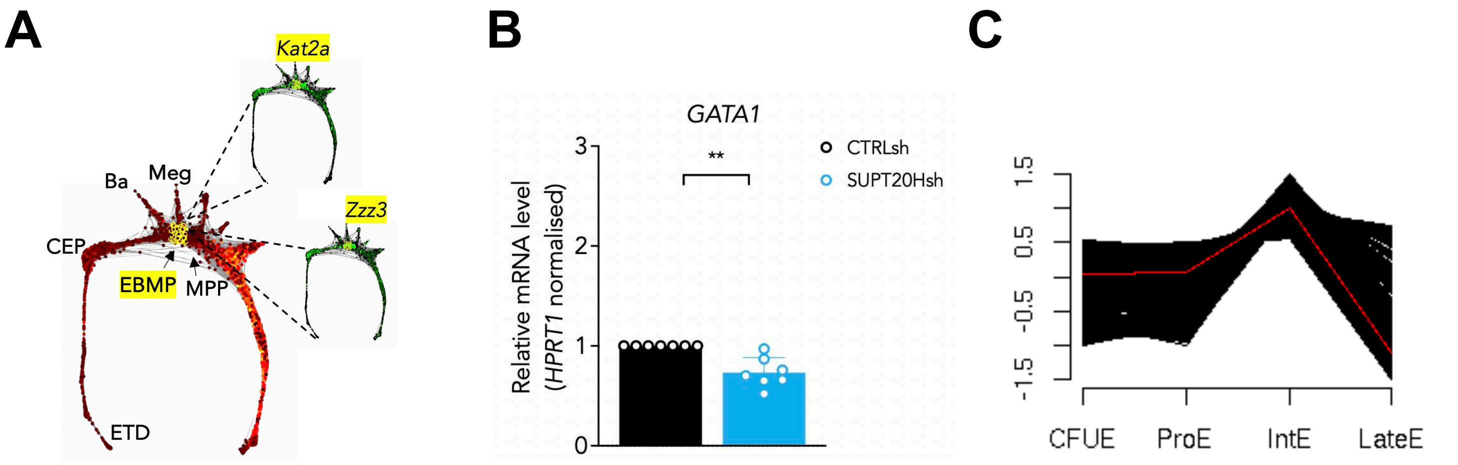


**Figure S5. Elements of SAGA associate with late erythroid specification programs. (relates to Figure 5)**
**(A)** Visualisation of gene expression (selected region) pattern of *Kat2a* and *Zzz3* using SPRING tool (https://kleintools.hms.harvard.edu/paper_websites/tusi_et_al/). As per Supplemental File 3, other genes in this region include regulators of E/Meg cell fate commitment including *Gata2*, *Zfpm1* and *Myb*. MPP: multipotent progenitors. EBMP: erythroid-basophil-megakaryocyte-biased progenitors. Meg: megakaryocyte. Ba: basophil. CEP: committed erythroid progenitors. ETD: erythroid terminal differentiation.
**(B)** Quantitative RT-PCR analysis of *GATA1* expression in SUPT20Hsh K562 cells. Mean ± SEM of 7 individual experiments; gene expression relative to CTRLsh, normalised to *HPRT1* housekeeping gene. Paired two-tailed t-test for significance **p<0.01.
**(C)** SAGA-specific elements peak at the Intermediate (IntE) phase of late erythroid differentiation. Representation of Cluster 17 obtained from the Human Erythroblast Maturation database (https://cellline.molbiol.ox.ac.uk/eryth/index.html). Details of individual genes can be found in Supplemental File 4.
